# Venom-inspired somatostatin receptor 4 (SSTR4) agonists as new drug leads for peripheral pain conditions

**DOI:** 10.1101/2024.04.29.591104

**Authors:** Walden E. Bjørn-Yoshimoto, Iris Bea L. Ramiro, Thomas Lund Koch, Ebbe Engholm, Ho Yan Yeung, Kasper K. Sørensen, Carolyn M. Goddard, Kathrine L. Jensen, Nicholas A. Smith, Laurent F. Martin, Brian J. Smith, Kenneth L. Madsen, Knud J. Jensen, Amol Patwardhan, Helena Safavi-Hemami

**Affiliations:** Department of Biomedical Sciences, University of Copenhagen; Copenhagen-N, Denmark; Department of Biochemistry, University of Utah; Salt Lake City, UT, USA; Department of Chemistry, University of Copenhagen; Frederiksberg, Denmark; Department of Neuroscience, University of Copenhagen; Copenhagen-N, Denmark; La Trobe Institute for Molecular Science, La Trobe University; Melbourne, Australia; Department of Anesthesiology and Pharmacology, University of Arizona; Tucson, AZ, USA; Department of Anesthesiology and Pain Management, Peter O’Donnell Jr. Brain Institute, UT Southwestern Medical Center; Dallas, Texas, USA; School of Biological Sciences, University of Utah; Salt Lake City, UT, USA

## Abstract

Persistent pain affects one in five people worldwide, often with severely debilitating consequences. Current treatment options, which can be effective for mild or acute pain, are ill-suited for moderate-to-severe persistent pain, resulting in an urgent need for new therapeutics. In recent years, the somatostatin receptor 4 (SSTR_4_), which is expressed in sensory neurons of the peripheral nervous system, has emerged as a promising target for pain relief. However, the presence of several closely related receptors with similar ligand-binding surfaces complicates the design of receptor-specific agonists. In this study, we report the discovery of a potent and selective SSTR_4_ peptide, consomatin Fj1, derived from extensive venom gene datasets from marine cone snails. Consomatin Fj1 is a mimetic of the endogenous hormone somatostatin and contains a minimized binding motif that provides stability and drives peptide selectivity. Peripheral administration of synthetic consomatin Fj1 provided analgesia in mouse models of postoperative and neuropathic pain. Using structure-activity studies, we designed and functionally evaluated several Fj1 analogs, resulting in compounds with improved potency and selectivity. Our findings present a novel avenue for addressing persistent pain through the design of venom-inspired SSTR_4_-selective pain therapeutics.

**One Sentence Summary:** Venom peptides from predatory marine mollusks provide new leads for treating peripheral pain conditions through a non-opioid target.

## INTRODUCTION

Persistent pain, also referred to as chronic pain, poses a significant global challenge, impacting more than one in five individuals and exacting substantial personal, societal, and economic burdens (*1*). Current therapeutic approaches depend on the type of pain and involve medications such as acetaminophen (paracetamol), non-steroidal anti-inflammatory drugs (NSAIDs), anticonvulsants, antidepressants, and opioids (*2*). Although generally effective for acute pain, these interventions often fail to sufficiently address severe persistent pain and can be associated with serious dose-limiting side effects, tolerance, and dependence, especially after prolonged use (*3*). In particular, the misuse of opioids has resulted in an epidemic of addiction and abuse of unprecedented scale (*4*), underscoring the urgent need for novel therapeutics and targets that act through opioid-independent pathways.

One such promising alternative target is the somatostatin receptor 4 (SSTR_4_), a member of the G protein-coupled receptor (GPCR) family activated by the peptide hormones somatostatin (SST) and cortistatin. The human SSTR family comprises five receptor subtypes (SSTR_1-5_) that share 46–63 % sequence identity and similar ligand-binding surfaces. Based on their similarities and evolutionary origin, the five subtypes can be divided into two groups: Group 1 comprises the SSTR_2,3_ and SSTR_5_ with sequence identities ranging from 50–58 %, and group 2 comprises the SSTR_1_ and SSTR_4_, sharing 63 % sequence identity (*5*). Activation of SSTRs, particularly group 1 SSTRs, is associated with inhibition of the secretion of various hormones (*6*). Different members of the SSTR family share the same endogenous ligands and are coupled to the Gα_i/o_ family, but have distinct expression profiles that can be associated with different physiological effects. For example, the SSTR_2_ and SSTR_5_ are highly expressed in neuroendocrine neoplasms, where their activation inhibits mitogenic signaling and growth (*6*). Somatostatin-based drug agonists that specifically activate these subtypes have long been used for the treatment of neuroendocrine disorders (e.g., acromegaly) and as diagnostic and therapeutic agents for certain types of cancers (*7-9*).

The first indication of the potential role of SSTR_4_-selective agonists for the treatment of pain came from the SSTR_1,4_-targeting peptide, TT-232, originally developed for the treatment of cancer but was later found to exhibit anti-inflammatory and anti-nociceptive effects (*10-12*). These results, as well as additional studies on the small molecule agonist J-2156 (*13, 14*), established the SSTR_4_ as a novel target for pain relief. Although the molecular mechanisms of SSTR_4_-mediated antinociception are not fully understood, several studies have suggested that SSTR_4_ activation in nociceptive neurons of the dorsal root ganglia (DRG) and trigeminal ganglia leads to downstream Gβγ-mediated activation of G protein-coupled inwardly rectifying potassium channels (GIRKs), allowing an outward flux of potassium ions, thereby hyperpolarizing cells involved in pain sensing and/or transmission (*15*). Furthermore, SSTR_4_ activation can reduce capsaicin-induced transient receptor potential cation channel subfamily V member 1 (TRPV1) currents in DRG neurons through Gα_i_ signaling, further hyperpolarizing the cell (*15*). Thus, SSTR_4_-selective ligands provide analgesia through the inhibition of peripheral pain signals and do not require central nervous system (CNS) administration or penetration.

Our recent discovery of a toxin agonist of group 2 SSTRs from a venomous cone snail, consomatin Ro1 (*16*), suggested that cone snail venoms may provide a unique source for the discovery of novel SSTR_4_ ligands. Consomatin Ro1 provided analgesia in two mouse models of pain, but only exhibited micromolar potency at the SSTR_4_ and was equipotent at the SSTR_1_ (*16*). To investigate whether cone snails evolved additional SSTR_4_-targeting toxins with improved potency and selectivity profiles, we computationally searched the venom gene datasets of 247 species, resulting in the identification of 529 SST-like lead sequences. Computational selection, synthesis, and receptor profiling of eight candidate toxins from this dataset led to the identification of consomatin Fj1, a potent and selective SSTR_4_ agonist that provides analgesia in postoperative and neuropathic pain models. Structure-activity studies using molecular dynamics (MD) simulations in combination with analog design revealed the peptide’s minimized binding mode and opportunities for further improvement of Fj1 for drug design and development.

## RESULTS

### Identification of candidate SSTR agonists from large venom gene library

We recently showed that cone snails have evolved peptide toxins that share sequence similarity with SST and its related peptides, cortistatin, urotensin II, and urotensin-related peptide (*17*). These toxins, referred to as “consomatins”, have evolved from an SST-like peptide used for endogenous signaling in cone snails. Following recruitment into the venom, these peptides greatly diversified to form a large family of SST-like toxins (*17*). To facilitate the selection of consomatin candidates that are likely to activate vertebrate SSTRs, with potential selectivity for the SSTR_4_, we extracted consomatin sequences from exon capture datasets of 247 cons snail species. We identified consomatin sequences in 169 species, in which we identified 529 sequences which subjected to principal component analysis (PCA) (see Methods for details). In addition to exon-capture data, consomatin Ro1 and consomatin Ro2, which were previously identified by transcriptome sequencing and mass spectrometric analysis of the venom of *Conus rolani*, as well as consomatin pG1 (previously named G1) from the transcriptome of *C. geographus*, were also included in the PCA (*16*). According to their prey, cone snails can be classified into worm hunters, snail hunters, and fish hunters. To enable comparative analysis of consomatins to endogenous SST-like hormones, peptide sequences from vertebrates (27 sequences), annelid worms (29 sequences), and mollusks (18 sequences) were included in the analysis. Consistent with our previous findings (*17*), toxins from worm hunting cone snails closely grouped with endogenous SST-like sequences from annelid worms, whereas many of the consomatins from fish hunters, including consomatin Ro1, grouped with vertebrate peptides (**Fig. 1A**). Consomatins were absent from snail hunting species. For the selection of toxin sequences from the PCA plot for chemical synthesis and SSTR receptor profiling, sequences from known fish hunting species were prioritized, as the SSTRs present in fish were highly similar to their human orthologs. Of these, sequences that grouped with vertebrate SST and related peptides were selected as these are more likely to have activity at the human receptors. The final list of sequences and their respective positions on the PCA is shown in **Fig. 1A-B)**.

**Fig. 1.**
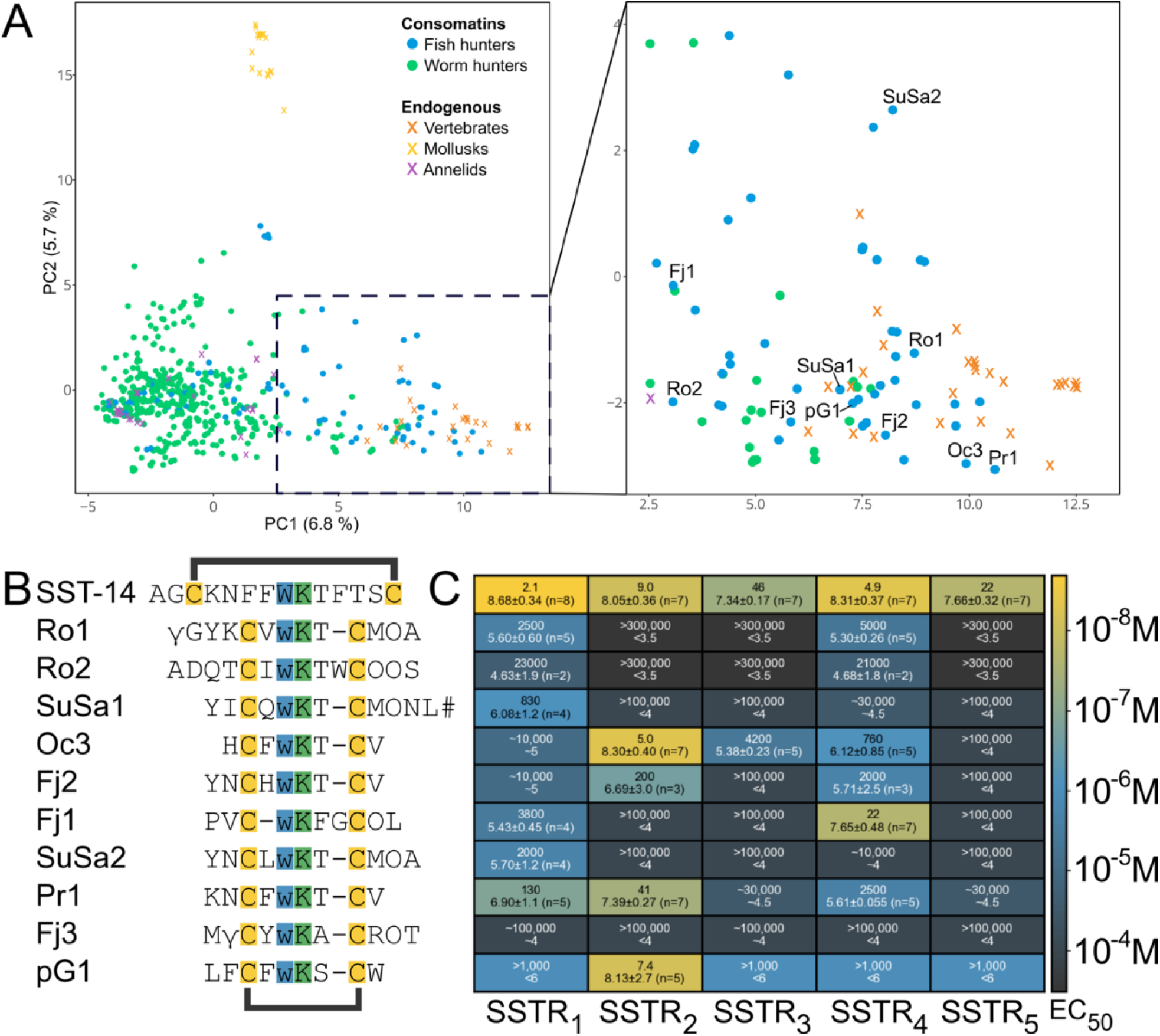
Computational selection and SSTR profiling of consomatins from a large toxin gene dataset. (**A**) Principal component analysis (PCA) of 529 SST-like toxin sequences was used as the basis for selecting lead compounds for SSTR screening. Percentages refer to the proportion of the total variance in the data explained by the given principal component. A subset of the PCA plot, including vertebrate SS-like sequences and selected consomatins, is shown (right box). (**B**) Synthesized consomatin sequences selected from the PCA plot. Cleavage sites and posttranslational modifications, such as D-Trp, were predicted based on the original somatostatin-like venom peptide, consomatin Ro1. Cysteines forming intramolecular disulfide bonds are shown in yellow and the essential Trp-Lys motif is highlighted in blue and green. Post-translational modifications are indicated by γ = γ-carboxyglutamate, w = D-Trp, O = hydroxyproline, and # = C-terminal amidation. SST-14 is human somatostatin-14. Seven of the listed sequences are from snails of the *Asprella* clade; Ro1 and Ro2 from *C. rolani*, Fj1, Fj2, and Fj3 from *C. fijisulcatus*, and SuSa1 and SuSa2 from *C. sulcatus samiae*, two are from the snails of the *Phasmoconus* clade; Oc3 from *C. ochroleucus*; Pr1 from *C. parius*, while pG1 was predicted from the transcriptome of *C. geographus* from the Gastridium clade (*16*). (**C**) Heatmap of activity values of selected consomatins tested at the five human SSTRs using the PRESTO-Tango β-arrestin recruitment assay. Rows correspond to the sequences shown in (B). Data are represented as EC_50_ (top line, nanomolar, two significant digits shown) and pEC_50_ ± 95 % confidence intervals (CI95) with the number of independent repeats in parentheses (bottom line, two significant decimals shown). Approximate values (∼) or “higher than” (>) and “lower than” (<) are used when we did not obtain full curves within the concentrations tested (see **Fig S1** for representative curves).

### Receptor profiling identifies potent and selective agonist of the SSTR_4_

The predicted mature peptide sequences selected from the PCA plot were synthesized to > 90 % purity using standard solid-phase peptide synthesis and verified using reverse-phase high-performance liquid chromatography and mass spectrometry. Post-translational modifications were predicted based on modifications previously observed for the venom peptide consomatin Ro1 (*16*). Modifications included γ-carboxylation of Glu, hydroxylation of Pro (Hyp), L-to-D epimerization of a Trp positioned within the disulfide loop, and C-terminal amidation based on the presence of a Gly-Arg/Lys motif (*18*). Disulfide bonds were predicted based on the presence of two cysteine residues. We note that the predicted modifications and the proteolytic N- and C-terminal cleavage sites may differ from those found in the native toxins. Future proteomics studies on collected venom are needed to establish the exact chemical identity of the toxins studied here.

Synthetic peptides were screened at the five human SSTRs using the PRESTO-Tango β-arrestin recruitment assay (*19*) (**Fig 1C** and **Fig S1**). Human somatostatin-14 (SST-14) and the previously characterized peptides, consomatin Ro1 and pG1, were included for comparison. The eight new peptides that were tested showed a range of SSTR activity profiles demonstrating that cone snail venoms are a rich source for the discovery of novel ligands of the human SSTRs. For example, Fj2 activates the SSTR_2_ and SSTR_4_ with EC_50_ values of 0.20 µM and 2.0 µM, respectively, Pr1 activates the SSTR_1_ and SSTR_2_ with EC_50_ values of 130 nM and 41 nM, respectively, and Oc3 shows activity at SSTR_2_, SSTR_3_, and SSTR_4_ with EC_50_ values ranging from 5.0 nM to 4.2 µM. Interestingly, although the sequence of Ro2 is distinct from the previously characterized Ro1, the two peptides that are both derived from the same cone snail species, *Conus rolani*, have very similar activity profiles at the human SSTR_1_ and SSTR_4_. Notably, none of the peptides tested showed any significant activity at SSTR_5_, although it should be noted that the peptides were tested for agonism only; thus, we cannot rule out antagonist activity. Of particular interest to us was Fj1, which activated the SSTR_4_ with an EC_50_ value of 22 nM, showing 173-fold selectivity over SSTR_1_ (EC_50_ = 3.8 µM) and no measurable activity at the other three receptor subtypes when tested at concentrations up to 100 µM.

### Consomatin Fj1 is a somatostatin “evolog” with a minimized receptor binding motif

Consomatin Fj1 is a ten amino acid long cyclic peptide from *Conus fijisulcatus* that contains four residues inside a disulfide loop and two residues at both its N- and C-termini (Sequence: Pro1-Val2-Cys3-D-Trp4-Lys5-Phe6-Gly7-Cys8-Hyp9-Leu10, **Fig. 2A-B**). The minimized disulfide loop of Fj1 closely mirrors that of the SSTR_2_-selective somatostatin drug analogs, such as octreotide and lanreotide. Incorporation of a shortened disulfide loop and a D-Trp in these analogs significantly prolonged their half-lives compared to the native human hormone (*9*). In contrast to these analogs, which were developed using medicinal chemistry approaches, consomatin Fj1 and other consomatins evolved from an endogenous SST-like signaling peptide following the principles of natural selection. Thus, we refer to these naturally evolved analogs as evologs (*16*). Notably, despite these distinctively different approaches, the final products exhibit a remarkably high degree of similarity (**Fig. 2B)**. One notable difference that distinguishes consomatin Fj1 from other SST analogs and evologs is that the characteristic D-Trp-Lys motif essential for SSTR activation (*6*) resides immediately following the first cysteine residue (**Fig. 2B**). As discussed in more detail below, these differences partly contribute to the potency and selectivity of the peptide for the SSTR_4_.

**Fig. 2.**
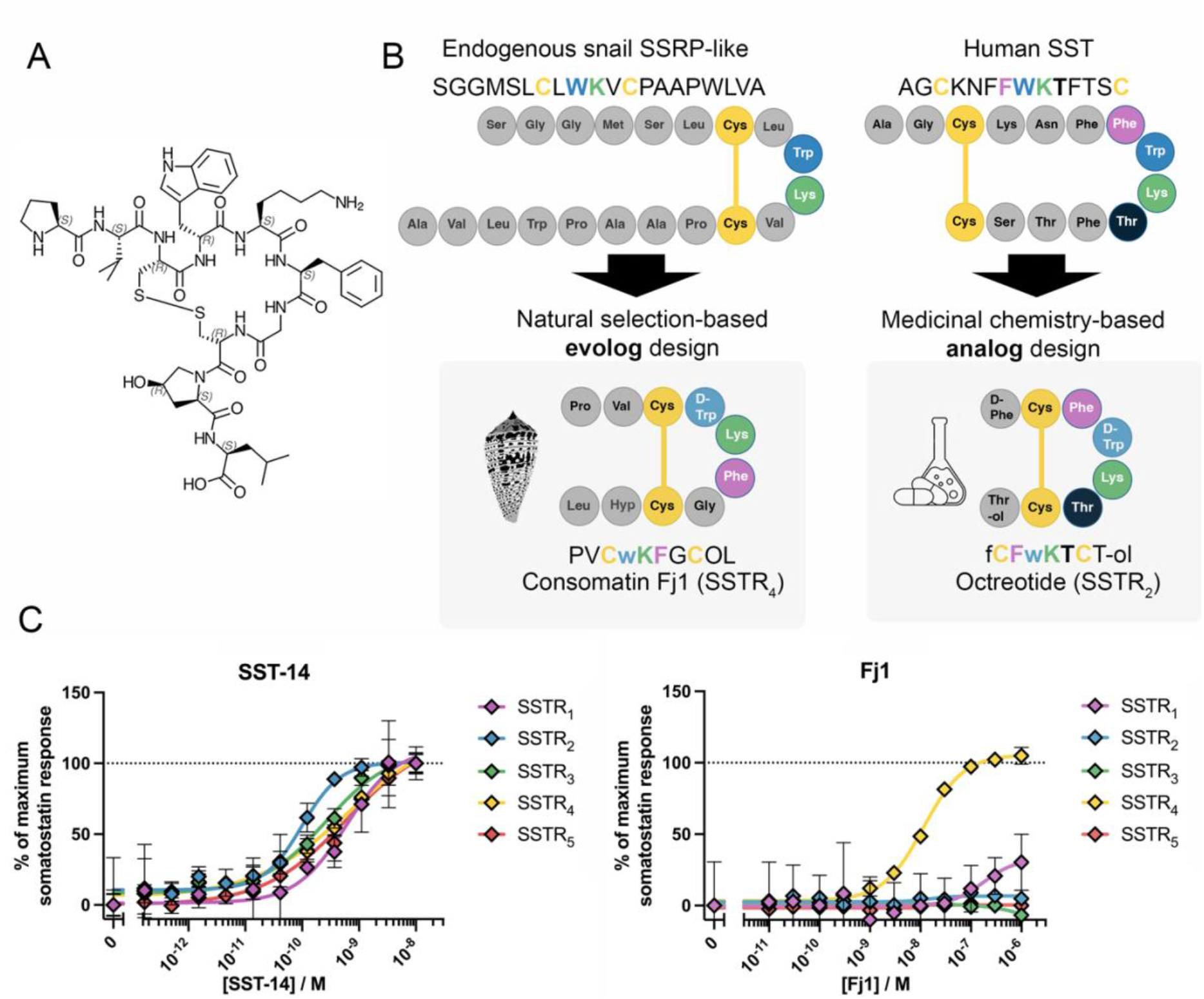
Consomatin Fj1 is a naturally evolved SSTR_4_-selective somatostatin evolog. (**A**) Chemical structure of consomatin Fj1, a ten-residue long cyclic peptide predicted from the venom gene-encoding exons of *Conus fijisulcatus*. Standard amino acid one or three letter abbreviations are used, extended with O/Hyp = 4-hydroxyproline, w/D-Trp = D-tryptophan, and -ol = C-terminal alcohol group in place of a carboxylic acid group. (**B**) Schematic comparison of the contrasting origins of consomatin Fj1 and the SST drug analog octreotide, both representing minimized SST scaffolds. Fj1 is a minimized SSTR_4_-selective evolog that originated from an endogenous SS-like signaling gene, while the SSTR_2_-selective drug analog octreotide was designed using medicinal chemistry approaches. (**C**) Representative concentration-response curves for G protein (Gα_oA_) dissociation of SST-14 and Fj1 at the five human SSTRs. Fj1 shows nM potency at the human SSTR_4_, with low potency at SSTR_1_ and no discernable activity at the other SSTR subtypes at 1 µM. Error bars represent the standard deviation of two technical replicates.

### Consomatin Fj1 induces G protein dissociation at the SSTR_4_

The analgesic effect of SSTR_4_ activation has been proposed to occur via GIRKs (*15*), and is largely mediated by the Gβγ subunits released from pertussis toxin-sensitive Gα_i/o_ proteins (*20*). Therefore, we assessed the activity of Fj1 at the SSTR_4_ using a bioluminescence resonance energy transfer (BRET)-based assay that measures GPCR activation by monitoring the association of labeled Gβγ subunits and a G protein-coupled receptor kinase (GRK) fragment, following dissociation of Gβγ subunits from the Gα subunit (*21*). We tested the activity of Fj1 at the SSTR_4_ using a range of different Gα proteins in the Gα_i_ family, including Gα_i1_, Gα_i2_, Gα_i3_, as well as the two dominant splice variants of Gα_o_, and observed similar potencies for each Gα protein (**Fig S2**). As the Gα_o_ type is the most highly expressed type in DRGs (*22*), the proposed site for SSTR_4_-mediated analgesia, we performed subsequent tests with the canonical Gα_o_ sequence (Gα_oA_). Here, we observed similar overall receptor activation profiles as in the PRESTO-Tango screening assay: potent activation of SSTR_4_ (EC_50_ = 6.0 nM) with only limited activity at SSTR_1_ (∼30 % activation at 1 µM) and no activation of the SSTR_2,3,5_ when tested at up to 1 µM (**Fig 2C**).

### Peripheral consomatin Fj1 administration provides analgesia in a post-operative pain model

Having established that consomatin Fj1 selectively activates the SSTR_4_ in the PRESTO-Tango and G protein dissociation assays, we next performed a dose response study for potential analgesic action in a model of post-operative pain, the paw incision model (*23*). C57BL/6J male mice injected intraperitoneally (i.p.) with 0.04–2.5 mg/kg of Fj1 and its effect on post-incision mechanical hypersensitivity was evaluated. Fj1 dose-dependently reduced post-incisional mechanical hypersensitivity, as measured by paw withdrawal thresholds in response to von Frey filament stimulation, with maximal effect observed at the dose of 2.5 mg/kg, and a peak effect observed at 1 h. The positive control used was morphine sulfate (5 mg/kg). While morphine exhibited higher peak efficacy than Fj1, the duration of action was shorter. Even at the 5 h time point, the mice treated with 2.5 mg/kg of Fj1 showed an increased paw withdrawal threshold of about 28 % of maximum Fj1 response (P = 0.0284, as compared to vehicle). Both in the groups treated with 0.04 mg/kg and with 0.1 mg/kg of Fj1, there was a similar trend as for 2.5 mg/kg, though neither reached a statistically significant difference from the vehicle control at any time point (P = 0.2936 and P = 0.0565, respectively, at 2 h).

### Peripheral consomatin Fj1 administration provides analgesia in a neuropathic pain model

Activation of the SSTR_4_ has previously been shown to provide analgesia in rodent models of neuropathic pain (*14, 24-26*). Drugs that are effective in post-operative pain may not be efficacious in neuropathic pain states and vice a versa. In general, neuropathic pain is particularly difficult to treat; many current treatment options are only effective in a limited subset of patients and can elicit severe side effects (*27-29*). To assess whether Fj1 could alleviate the mechanical hypersensitivity associated with neuropathic pain, the peptide was tested in the spared nerve injury (SNI) model of peripheral neuropathic pain using male C57BL/6 mice. The sciatic nerve was injured on one side of the mouse (ipsilateral side), whereas the other side (contralateral side) served as a control. On day 16 post-injury, either 0.5 mg/kg or 5 mg/kg of Fj1, 30 mg/kg of gabapentin, or vehicle was administered by i.p. injection (**Fig. 3B**). Both doses of Fj1 reduced mechanical hypersensitivity 30 min after injection, and this effect was more pronounced after 1 h. For the 0.5 mg/kg dose, the effect persisted at 2.5 h, while for the 5 mg/kg dose, the tendency was the same, albeit not reaching statistical significance (P = 0.0548 compared to vehicle). These results demonstrate that Fj1 can provide potent and efficacious analgesia in this model, although with a slower onset and shorter duration of action than 30 mg/kg of gabapentin.

**Fig. 3.**
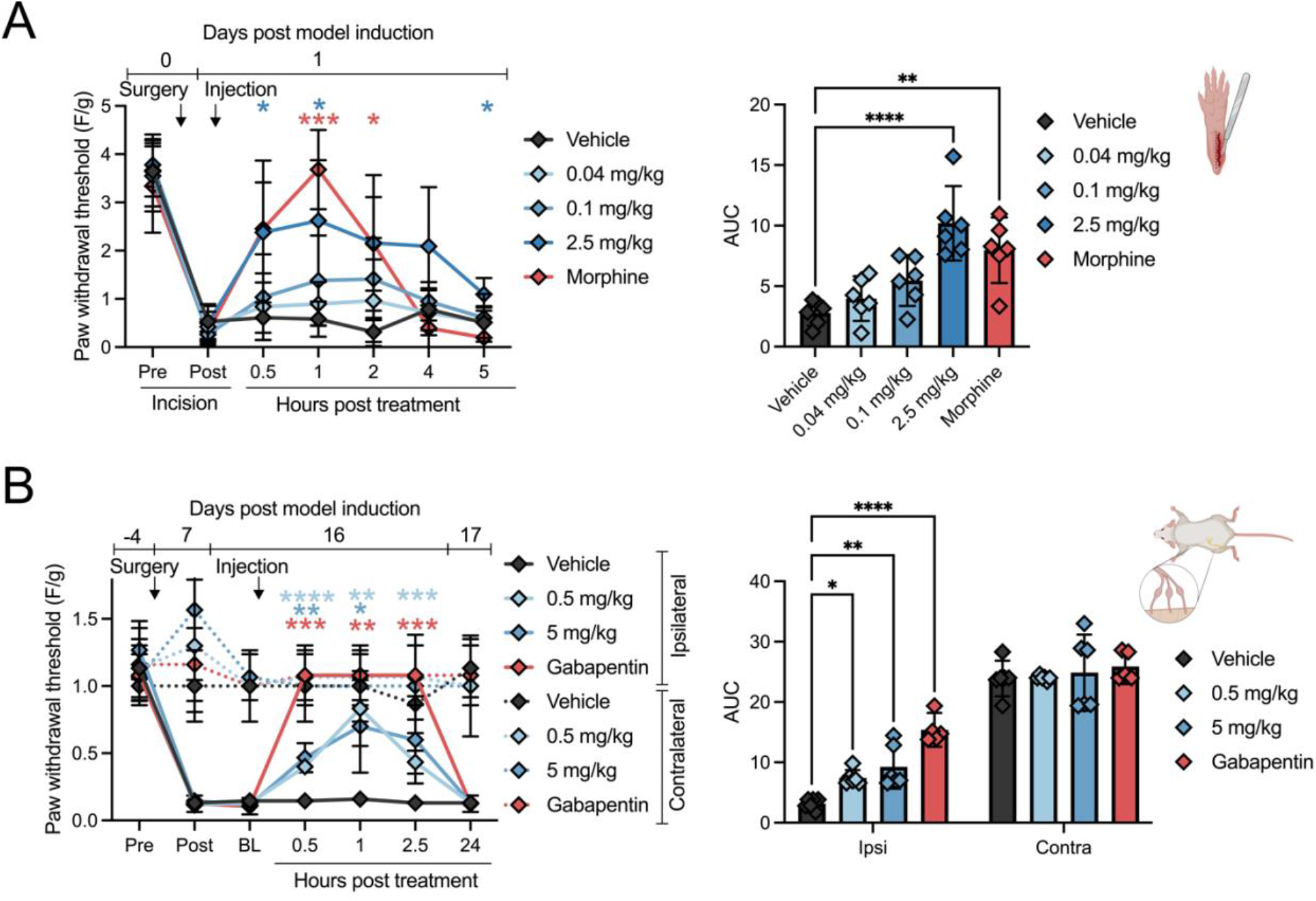
Consomatin Fj1 provided analgesia in two mouse models of pain. (**A**) Post-surgical pain model (paw incision). Plantar incision surgery was performed 24 h post-surgery, and mice were administered morphine sulfate (5 mg/kg), indicated doses of Fj1, or vehicle (normal saline) by intraperitoneal (i.p.) injections, whereafter mechanical hypersensitivity was evaluated at indicated times (left). Values represent mean ± CI95 from six mice. A summary of the area under the curve is shown (right), where each data point represents one mouse, and the error bars represent CI95. (**B**) Neuropathic pain model (spared nerve injury, SNI). The peroneal and tibial branches of the sciatic nerve of the left hindleg were ligated, 16 days post-surgery, the mice were administered gabapentin (30 mg/kg), indicated doses of Fj1, or vehicle by i.p. injections, whereafter mechanical hypersensitivity was evaluated at indicated times. Values represent the mean ± CI95 from five (gabapentin) or six (all other conditions) mice. Statistical analysis was performed as described in the methods section. Asterisks indicate P-values < 0.05 (^*^), 0.01 (^**^), 0.001 (^***^), and 0.0001 (^****^) as compared with the vehicle control.

### Molecular dynamics simulation reveals the binding mode of Fj1 at the SSTR_4_

To investigate the binding mode of Fj1 at the SSTR_4_ and better understand the molecular basis for the peptide’s selectivity and potency, we performed molecular dynamics (MD) simulations in the µs range of Fj1 to the holo-SSTR_4_ embedded within a 1-palmitoyl-2-oleoyl-*sn*-glycero-3-phosphocholine (POPC) bilayer. The binding mode of consomatin Fj1 was stabilized and almost exclusively characterized through the side chain interactions of D-Trp4 and Lys5 with receptor residues (**Fig. 4A**). D-Trp4 buried deep within the hydrophobic core cavity formed by the side chains of Leu123^3×29^, Met130^3×36^, Phe131^3×37^, Ile181^4×61^, Leu200^45×52^, and Phe275^6×51^ (superscripts denote the GPCRdb numbering scheme outlined by Isberg *et al*. (*30*)). Interestingly, D-Trp4 occupied the same binding pocket as the corresponding L-Trp of SST-14 observed in the recent cryogenic electron microscopy (cryoEM) structure of SSTR_4_ (PBD ID: 7XMS) (*31*) (**Fig. 4B**). Lys5 forms a salient salt bridge with the side chain of Asp126^3×32^, also mirroring the conformation and interactions of the corresponding Lys9 in SST-14. Additionally, Lys5 in Fj1 also demonstrated transient yet notable hydrogen bonding to the side chain of Ser300^7×41^ during simulation. On the opposite face of the side chain, Lys5 maintains a side-by-side stacking arrangement against the indole ring of D-Trp4, which is also present in the monomeric structure of Fj1, and is characteristic of SST receptor binding and activation (*6*). Both D-Trp4 and Lys5 demonstrated limited sampling of other conformations throughout the simulation.

**Fig. 4.**
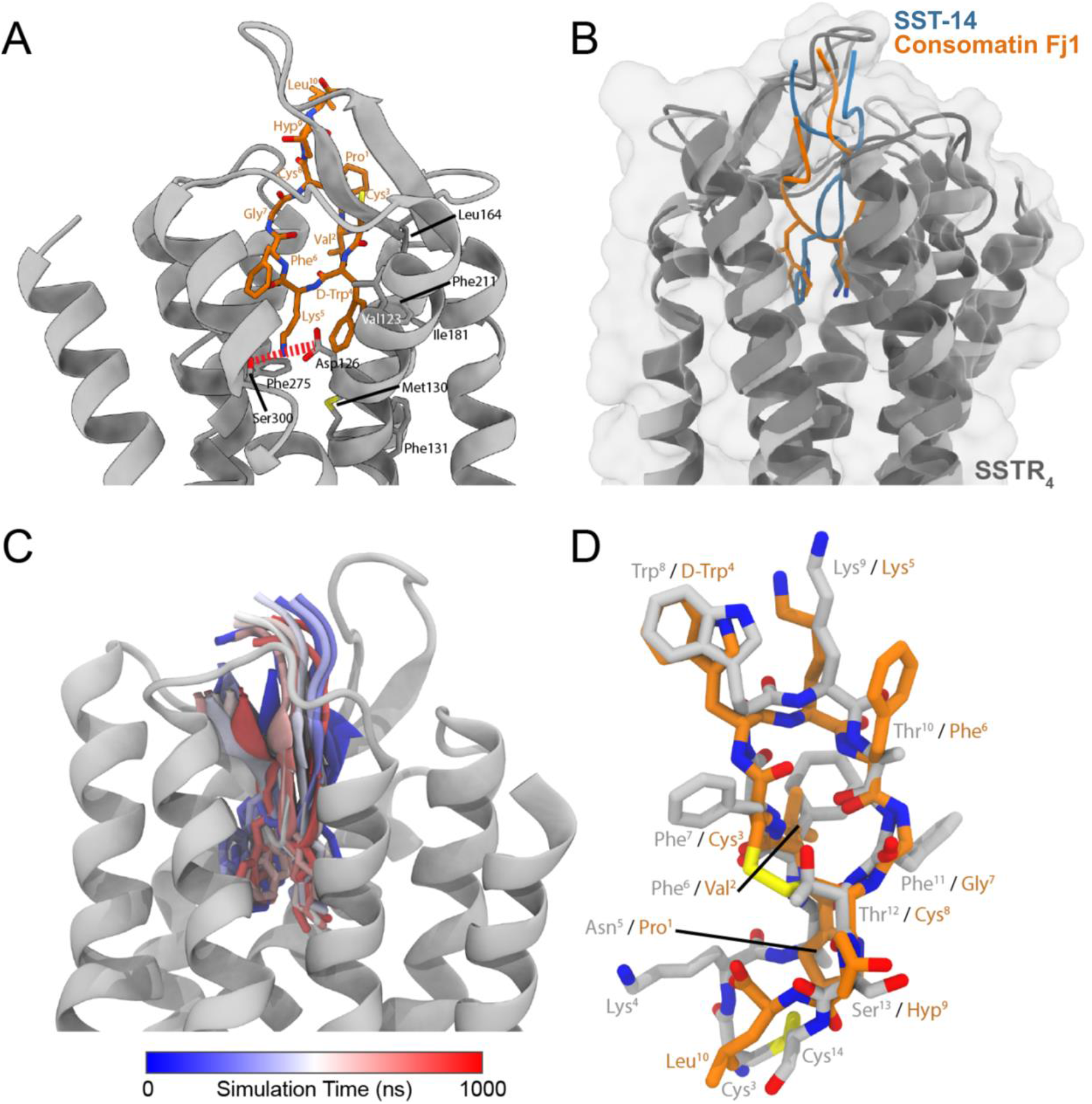
Molecular Dynamics (MD) simulations reveal contacts between Fj1 and SSTR_4_. (**A**) Representative orientation of Fj1 (orange stick representation) in complex with SSTR_4_ (gray ribbon representation). The membrane present within the simulation was removed for clarity. The interactions made by Fj1, which facilitated binding, are labelled. The representative binding mode was obtained by clustering all replicate trajectories, with a model representing the largest cluster displayed. (**B**) Overlay of Fj1 (orange) and SST-14 (blue, PDB 7XMS) in complex with SSTR_4_ (dark gray). (**C**) The orientation of Fj1 throughout a representative 1 µs simulation is shown with 10 snapshots colored from blue to white to red. (**D**) Representative orientation of Fj1 (orange) from the simulation when in complex with SSTR_4_, overlaid with SST-14 in complex with the same receptor construct (gray, PDB 7XMS).

In contrast to the persistent orientation adopted by D-Trp4 and Lys5, residues bordering and outside the cyclic core, Pro1, Cys3, Phe6, Cys8, Hyp9, and Leu10 (i.e., with the exception of Val2) displayed considerable mobility, demonstrating limited persistent contacts describing the ligand-bound orientation (**Fig. 4C-D**). Likewise, residues outside of the conserved pharmacophore (Phe7-Trp8-Lys9-Thr10) in SST-14 do not form significant side chain interactions with SSTR_4_, as has been observed with SSTR_2_ (*31*), instead forming more backbone and intramolecular interactions (most notably between Phe6-Phe11). The short cyclic core (four residues in Fj1, compared to ten residues in SST-14) does not protrude significantly beyond the binding pocket and aligns well with the smaller four-residue pseudo-core formed between Phe6 and Phe11 of the SSTR-bound structure of SST-14. Val2 presents a somewhat smaller hydrophobic interface within the helical core, packing between residues Val212^5×40^ of helix 5 and Leu283^6×59^ of helix 6, occupying a position similar that to of Phe7 in SST-14. These simulations showed that Fj1 binds SSTR_4_ in a manner similar to SST-14, particularly in regard to the Trp-Lys motif, and that residues outside of this could be amenable to modification while retaining SSTR_4_ binding.

### Analog design provides improved agonists and scaffolds for future drug development

To assess the molecular determinants of SSTR_4_ potency and selectivity in the G protein dissociation assay, we first performed alanine scanning of the four amino acids in the loop that were buried in the binding pocket of the receptor (**Fig. 5**). As expected, mutating key residues known to be important for the binding of somatostatin (*6*), Trp4 (Fj1A1) and Lys5 (Fj1A2), to Ala markedly decreased potency by 5000-fold and 730-fold, respectively. The mutation of Phe6 (corresponding to Thr10 in SST-14, see **Fig. 4D**) to Ala (Fj1A3) also decreased the potency by 630-fold. However, mutating either Gly7 to an Ala (Fj1A4) or deleting it, thereby shortening the loop (Fj1A5), resulted in a slight increase in potency at SSTR_4_ (6.2-fold and 65-fold, respectively) with no discernable increase in potency at other SSTRs. Next, we assessed the role of the predicted posttranslational modifications. Changing D-Trp and L-Hyp, to their unmodified equivalents, L-Trp and L-Pro (Fj1A6 and Fj1A7, respectively), had virtually no effect on the SSTR_4_ potency, although both showed an increase in SSTR_1_ potency. Another epimerization, the substitution of L-Phe6 to D-Phe (Fj1A8), on the other hand decreased the potency at SSTR_4_ by 42-fold. Since the MD simulations suggested that residues outside the cyclic core did not form significant and salient interactions with the receptor, we tested the “minimal core” peptide Cys3-D-Trp4-Lys5-Phe6-Gly7-Cys8 (Fj1A9). This decreased the potency by 47-fold, while deleting only the N-terminus (Fj1A10) resulted in a 13-fold decrease in potency. However, deletion of only the C-terminus (Fj1A11) resulted in an analog that was virtually equipotent at the SSTR_4_.

**Fig. 5.**
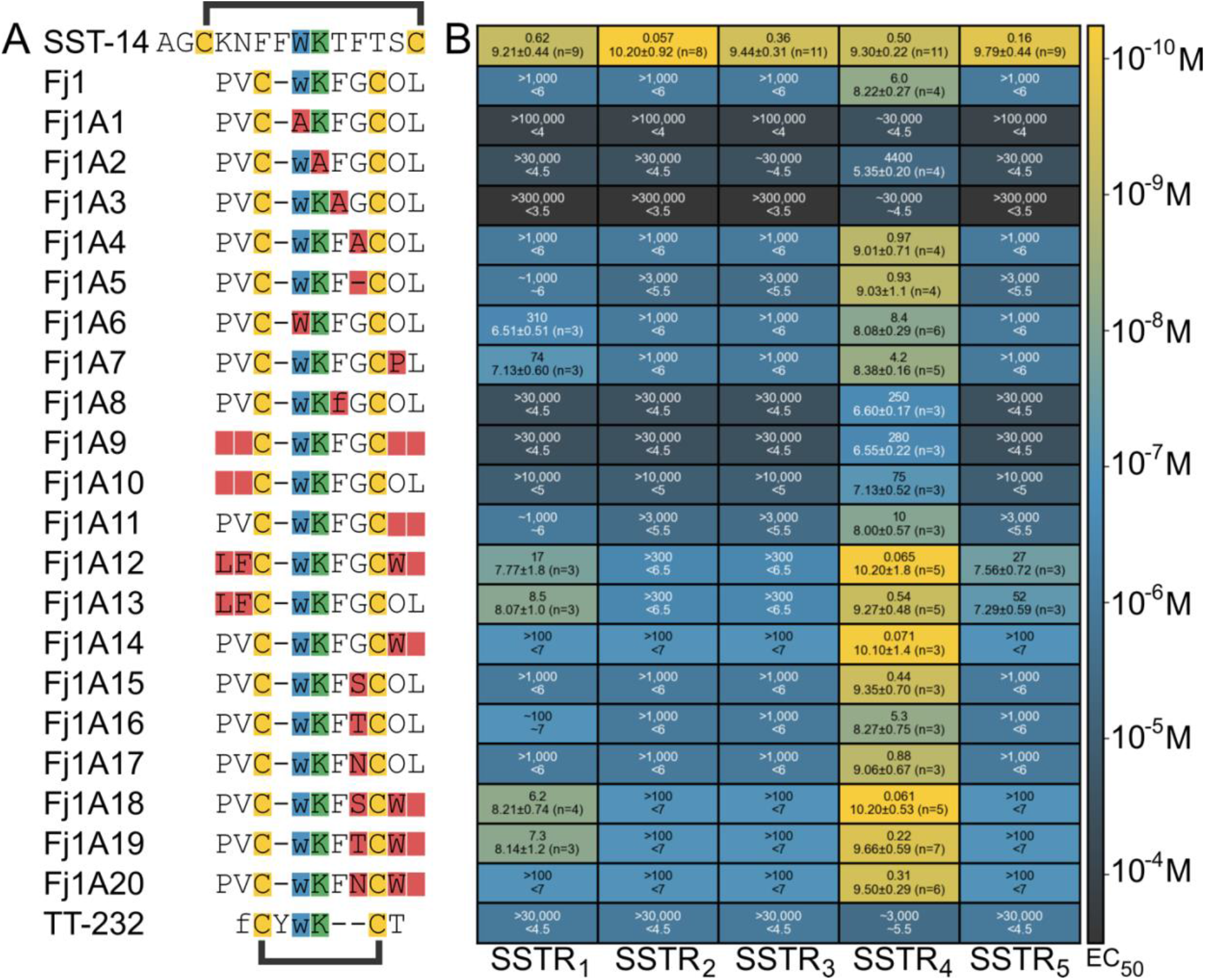
Analog design and testing identified improved ligands for SSTR_4_-selective drug development. (**A**) Identifiers and sequences of SST-14, Fj1, and the analogs of Fj1. Sequences (left) show the Cys residues forming the intramolecular disulfide in Fj1 and its analogs highlighted in yellow. For all sequences, the Trp residue is shown in blue, the Lys residue in green, and all modifications in the analogs, as compared to Fj1, are highlighted in red. Sequences use standard amino acid one-letter abbreviations extended with O=hydroxyproline and lower-case letters denoting D-amino acids. (**B**) Heatmap showing the potency of Fj1 and its analogs at the five human SSTRs in the G protein dissociation assay using Gα_oA_. The text in each cell represents the EC_50_ value (first line, nanomolar, two significant digits) and the pEC_50_ ± CI95 (two significant decimals) with the number of independent repeats for each in parentheses (second line). Approximate values (∼) or “higher than” (>) and “lower than” (<) are used when we did not obtain full curves within the concentrations tested (see **Fig S3** for representative curves). The known SSTR_1_ and SSTR_4_ peptide agonist, TT-232 was also tested. However, we note that we could not dissolve it at sufficiently high concentrations to fully test this peptide in the assay used).

Recently, we identified consomatin pG1 as a potent and selective agonist of the SSTR_2_ (*16*). To investigate whether determinants from pG1 could be used to improve the potency of Fj1, we designed a hybrid analog combining the cyclic core residues of Fj1 with the amino acids flanking the pG1 core (Fj1A12). This hybrid analog showed a 92-fold increase in potency at the SSTR_4_, although with a corresponding increase at the SSTR_1_ and SSTR_5_. Interestingly, this did not lead to increased potency at the SSTR_2_. Introducing only the N-terminus of pG1 (Fj1A13) increased potency by 11-fold at the SSTR_4_, with SSTR_1_ activity similar to that of Fj1A12. With the suggestion from Fj1A11, that the Fj1 C-terminus was not required, we next tested the effect of adding the C-terminus from pG1, namely the single Trp residue, to Fj1. This analog (Fj1A14) showed an 85-fold increase in potency at the SSTR_4_, with no enhanced activity observed at the other receptor subtypes, including the SSTR_1_. Given the apparent redundancy of Gly7, we evaluated whether replacing Gly7 with hydrophilic amino acids to decrease the hydrophobicity of the peptide could be tolerated. Substituting Gly9 with Ser (Fj1A15) or Asn (Fj1A17) slightly increased the potency at SSTR_4_ (14-fold and 6.8-fold, respectively), whereas a Thr at this position (Fj1A16) retained the SSTR_4_ potency of Fj1, while showing a slight increase in SSTR_1_ activity. Based on these observations, we next assessed whether the increased potency of Fj1A14 could be combined with the polar substitutions of Gly7. While a Ser in place of Gly7 (Fj1A18) was well tolerated at the SSTR_4_, both the Thr and Asn substitutions (Fj1A19 and Fj1A20, respectively) showed a slight decrease in SSTR_4_ potency compared to Fj1A14. However, all three double substitutions increased SSTR_1_ activity, thereby decreasing selectivity.

These results show that while Fj1 was the most potent and selective SSTR_4_ peptide agonist identified from venom-encoding genes, there are clear opportunities for further optimization of the peptide’s potency, selectivity, and physiochemical properties such as hydrophilicity.

## DISCUSSION

Cone snail venoms represent a rich source for the discovery of biomedical tools and drug leads. Among these, the cone snail peptide ω-MVIIA derived from the venom of the magician cone, *Conus magus*, is arguably the most prominent example (*32*). MVIIA, also known as ziconotide, inhibits pain signals by selectively blocking the N-type calcium channel, Ca_v_2.2, expressed in the CNS (*33*). Ziconotide not only revealed the existence of the Ca_v_2.2 channel, but also delineated its role in pain signaling and became an approved drug for the treatment of chronic intractable pain in 2004 (*34*). Another analgesic cone snail peptide toxin that modulates pain signals in the CNS and has proven analgesic effects in humans is contulakin-G, also known as CGX-1160, derived from the venom of the geographer cone, *Conus geographus* (*35*). Contulakin-G shares sequence similarity with the human neuropeptide, neurotensin. Although the exact mode of action of contulakin-G remains unknown, recent studies suggest that, *in vivo*, this peptide acts as an agonist of the neurotensin 2 receptor (NTSR2), leading to the inhibition of the R-type calcium channel, Ca_v_2.3, expressed in the spinal cord (*36*). Contulakin-G entered a phase Ia clinical study for the treatment of spinal cord injury related pain, one of the most difficult to treat type of pain conditions. Despite its significant effects in reducing pain, the clinical development of contulakin-G was discontinued when the company developing this peptide shut down (*34*).

Our recent discovery of consomatin Ro1, an analgesic toxin from *Conus rolani* that activates the human SSTR_1,4_, provided another lead compound for pain from cone snail venom. Unlike MVIIA and contulakin-G, consomatin Ro1 targets receptors expressed in the peripheral nervous system and has no analgesic effect when administered centrally in mice (*16*). This peripherally restricted site of action can be advantageous in two different ways. First, the delivery of the drug will not require invasive techniques such as intrathecal or epidural puncture. Next, the likelihood of CNS-related side effects such as dependence or respiratory depression are significantly reduced. Here, we describe the discovery and preclinical evaluation of another SSTR_4_ agonist with superior potency and selectivity, consomatin Fj1, derived from *Conus fijisulcatus*.

Based on their relatedness, cone snails can be grouped into approximately 44 distinct clades (*37*). Snails belonging to the same clade typically share similar venom compositions and hunting behaviors, although exceptions exist. Notably, *C. fijisulcatus* and *C. rolani* both belong to the *Asprella* clade, one of eight known lineages of fish hunting cone snails (*37*). Unlike most other cone snails that inhabit shallow coastal waters, *Asprella* snails live offshore at depths of 60 – 250 m (*38*). *Asprella* snails use an unusual hunting strategy characterized by venom injection followed by a slow onset of venom action. We previously observed predation by *Asprella* snails in captivity, which took between 15 min and 3 h from envenomation until the prey was incapacitated^16^. We hypothesized that this “ambush-and-assess” hunting strategy is accompanied by the evolution of toxins that inhibit the prey’s escape response. Furthermore, the slow onset of action suggested that the venom of *Asprella* snails may not be dominated by classical ion channel modulators but may contain toxins that target GPCRs. Here, to discover novel ligands of human SSTRs, we used PCA to identify consomatins that group with human SST and its related peptides and prioritized toxins that belonged to species of the *Asprella* clade. In the small set of *Asprella* consomatins we synthesized and tested, all had activity at the human SSTRs, with a notable bias towards activation of the SSTRs in group 2 (SSTR_1_ and SSTR_4_). Only a single *Asprella* consomatin showed notable activity at a group 1 SSTR at the concentrations tested (Fj2 at the SSTR_2_). There is emerging evidence for the role of agonists group 2 SSTRs in inhibiting pain and inflammation in rodent models of disease, as well as recent clinical data from a phase II clinical trial showing that an SSTR_4_ agonist is effective for treating peripheral diabetic neuropathy (*39*). Given the observed predominance of SSTR_1_ and SSTR_4_ activity of consomatins identified from only five *Asprella* species of at least 18 known (*40, 41*), there are likely to be more consomatins of interest for mammalian pain modulation in this clade of fish hunting cone snail (*42*).

Because of its high potency and selectivity for the SSTR_4_, we focused our subsequent structure-activity studies on consomatin Fj1. An unusual feature of consomatin Fj1 is its unique sequence motif, with the canonical Trp-Lys dyad immediately following the first Cys residue. To better understand the role of this dyad and the overall mode of action of the peptide, we performed MD simulations combined with functional studies on select analogs. MD simulations of Fj1 and comparison with the known structure of SST-14 bound to the SSTR_4_ revealed a similar binding mode between the toxin and the human peptide. Where SST-14 utilizes the two Phe residues in positions 6 and 11 to essentially create a T-shaped π-π interaction with Phe7, Trp8, Lys9, Thr10 in between, in Fj1, these aromatic residues are replaced with Cys residues, creating a covalent bond linking the corresponding residues, and retaining the four-residue “loop” that SST-14 transiently inhabits in its SSTR-bound form (see **Fig. 4B,D** as well as work by Robertson *et. al* and Zhao *et al*. (*31, 43*)). The D-Trp-Lys motif remained in the lower binding pocket in the Fj1 simulation, where residues interacted largely with the same SSTR_4_ side chains as the corresponding residues in SST-14. Notably, the D-Trp of Fj1 inhabits the same binding pocket as the corresponding L-Trp of SST-14, which likely explains why we observed little effect of changing D-Trp to L-Trp in (Fj1A6). While we did observe a slight difference in SSTR_1_ potency, the L-to-D epimerization that was previously observed in consomatin Ro1, and predicted in other consomatins, including Fj1, likely plays a larger role in stabilizing the minimized fold and providing resistance to proteolytic cleavage, as this modification is known to evade recognition by endogenous enzymes and has been introduced in other peptide drugs, including SST analogs (*44*). To identify the minimum binding motif at the SSTR_4_ we tested the “core” motif of Fj1 linked by the disulfide loop (Cys-D-Trp-Lys-Phe-Gly-Cys; Fj1A9). This analog retained both activity and selectivity, albeit being approximately 47-fold less potent at SSTR_4_. In line with this observation, residues outside this core did not show notable static side chain-side chain interactions with the SSTR_4_ throughout the simulation. However, analog screening suggested a differential role for the N- and C-terminal residues in binding. Removing the two N-terminal residues which, due to the “shifted” D-Trp-Lys motif in relation to the cysteines, are situated closer to the binding pocket surrounded by TM5 and TM6 (Fj1A10), showed a potency loss of 13-fold. In comparison, removing the two C-terminal residues (Fj1A11), which protrude from the binding pocket and are surrounded by the flexible ECL2 and ECL3 in the MD simulation, barely changed the potency at the SSTR_4_. As most of the polar interactions with residues outside the core of Fj1 involve backbone residues, these findings suggest the possibility of introducing additional modifications outside the core. Indeed, adding the C-terminal Trp from consomatin pG1, a potent and selective agonist of the SSTR_2_, to the Fj1 scaffold resulted in a significant increase in potency while retaining selectivity (Fj1A12). This surprising finding demonstrates that there is significant room for improving the potency of Fj1 at the human SSTR4 without compromising the selectivity of the peptide. Additionally, analog screening demonstrated that Gly7, which is located within the disulfide loop, is amenable to deletion (Fj1A5) and substitutions (Fj1A15-Fj1A20) that can retain or even improve potency while introducing hydrophilic side chains. However, some of these come at the expense of decreased selectivity for SSTR_4_ over SSTR_1_.

The small molecule SSTR_4_ agonist, J-2156, which has been reported to be about 360-fold selective for SSTR_4_ over SSTR_1_ and 390-fold over SSTR_5_ in binding assays (*45*), has been widely used in the literature (*14, 15, 24, 26, 45, 46*). However, peptide agonists that can distinguish between SSTRs in group 2 (SSTR_1_ and SSTR_4_) have not been previously reported. The most widely used peptide agonist at the SSTR_4_ is the heptapeptide, TT-232, which suffers from low potency and a 6.5-fold reported selectivity over SSTR_1_ (*47*). Using the venom peptide Fj1 as an inspiration, this study provides several analogs with significantly improved potency and selectivity over previously described peptide agonists and suggests additional opportunities for further optimization.

Moreover, having a potent and selective peptide agonist of the SSTR_4_ expands the repertoire of existing drug leads for this target from small molecules to peptides. Peptides, especially those that have evolved by nature, often display better selectivity and fewer off-target effects or complications from hepatic metabolism (*48*). While the *in vivo* half-life of peptides is often limited by rapid renal clearance, the design of peptide conjugates can significantly increase the duration of action, as seen with many recent peptide therapeutics that now have weekly, instead of daily, dosing regimens (*49*). The importance of these advantages is reflected by the rapidly increasing number of approved peptide therapeutics in recent years (*50, 51*). Here, we show that the small cyclic peptide Fj1 can efficiently alleviate the mechanical hypersensitivity associated with two types of pain in rodent models of the disease.

The incision model of postoperative pain gives rise to a complex pain response dependent on multiple components, sensitizing both C- and Aδ-fibers that innervate the area around the injury (*52*). The expression pattern of the SSTR_4_ in C- and Aδ-fibers is not well established, but differences here, along with the central effects of opioids, could help explain the difference in the maximal response observed between Fj1 and morphine. The expression of SSTR_4_ is upregulated in inflammatory models (*53*), and activation can inhibit inflammation induced by either mustard oil or lipopolysaccharide (*10, 54, 55*). In response to injury, inflammatory responses are generally observed within 2 h (*56, 57*). SSTR_4_-mediated inhibition of inflammation could help explain the seemingly longer-lasting effect of Fj1 compared to morphine, where the later stage effect might, in part, be due to a reduction in the post-injury inflammatory response that would otherwise hypersensitize the pain response (*58*). However, it is also possible that the effects are due to a slower off-rate at the receptor, or simple pharmacokinetic effects. A thorough examination of these effects is outside the scope of this study.

Similar to the effect of Fj1 in the paw incision model, the maximum observed effect of a single i.p. dose of Fj1 in the SNI model was also observed at the 1 h time point for both doses used. At this time point, the 5.0 mg/kg dose did not elicit a larger effect than the 0.5 mg/kg, suggesting that 0.5 mg/kg was a saturating dose in this model. At the 2.5 h timepoint there was a tendency for a more pronounced effect for the higher dose, which is likely a pharmacokinetic effect reflecting clearance. Our preliminary investigations into the analgesic effects of Fj1 demonstrated that this peptide can effectively alleviate pain like behavior in two mechanistically distinct mouse models. Future studies on Fj1 and its analogs in additional pain models and species will provide insights into the versatility of these new SSTR_4_ agonists and likely inform on the design of additional analogs with improved *in vivo* potency and efficacy profiles.

Our study has several limitations. While the MD simulations of Fj1 confirmed our experimental data and supported or analog screen, an experimental structure is needed to further verify the exact nature of the binding determinants. Furthermore, although we report various potent and selective SSTR4 agonist these compounds will likely require optimization of their pharmacokinetic profiles before advancing to clinical approval. Finally, while Fj1 and other SSTR4 agonist are capable of alleviating pain in rodent models, clinical validation of the SSTR_4_ is incomplete. Recent phase 2 clinical trials on a small molecule SSTR_4_ agonist provided preliminary target validation for diabetic neuropathy, while trials for osteoarthritis and chronic lower back pain showed no effect over placebo (*39*). No clinical trials for therapeutics targeting the SSTR_4_ for postoperative pain have been done. As such, which animal models will ultimately translate to humans when targeting the SSTR_4_ is yet unknown.

In conclusion, the discovery of Fj1 and other cone snail peptides with distinct activity profiles at the human SSTRs expands our knowledge of the chemical and pharmacological diversity of the compounds evolved by these marine predators. Given the diversity of somatostatin evologs identified, we strongly anticipate that future functional characterization of these peptides will continue to provide a stream of novel ligands of the SSTRs and potentially other GPCRs with unique selectivity and potency profiles. Our findings regarding the broad use of somatostatin evologs in diverse cone snail species further suggest the existence of similar peptides in other venomous animals. Finally, the apparent richness of group 2 SSTR-targeting peptides in *Asprella* snails highlights the usefulness of this clade in the discovery of novel SSTR_4_-targeting analgesic leads.

## MATERIALS AND METHODS

### Study design

This study consists of preclinical call-based and rodent models, as well as computational approaches, designed to investigate the potential for identifying lead compounds targeting the human somatostatin system in the venom of marine cone snails, as well as the suitability of an identified peptide as a lead for postoperative and neuropathic pain conditions. The number of replicates and repeats are indicated and all statistical tests, curve fitting, and other data analysis are described in the Materials and Methods section. All animals randomly assigned to their respective treatment groups animal and the experimenters performing measurements were blinded to the treatment. The number of animals was chosen by the group performing the experiments, based on their experience, to ensure sufficient statistical power.

### G protein dissociation assay

HEK293 cells (ATCC) maintained in DMEM (Gibco) supplemented with 10% FBS (Biowes, Nuaillé, France) and 100 U/mL penicillin / 100 mg/mL streptomycin (Gibco) (growth medium) were maintained in a water jacketed 5 % CO_2_ incubator and passaged every 2-3 days using trypsin/EDTA (Gibco). 1E6 cells were seeded into 6 well plates. The following day, the medium was changed, and the cells transfected by adding 0.33 µg of SSTR construct, 0.66 µg of Gα protein, 0.33 µg Venus 156-239-Gβ, 0.33 µg Venus 1-155-Gγ, and 0.33 µg masGRK3ct-Nluc DNA into 27 µL of 150 mM NaCl, and 12 µL of 100 mg/mL linear polyethyleneimine (PEI) at molecular weight 25 kDa (Polysciences, Warrington, PA, USA) into 38 µL of a 150 mM NaCl solution. The two tubes were mixed and incubated for 10-15 min before addition to the cells. The following day, 5E4 cells per well were seeded in 100 µL of growth medium into poly-D-lysine-coated white 96 well OptiPlates (PerkinElmer). The following day, the cells were washed in 80 µL of HBSS supplemented with 20 mM HEPES, 1 mM CaCl_2_, and 1 mM mgCl_2_ (assay buffer), and 80 µL of assay buffer supplemented with 0.1 % BSA (stimulation buffer) was added. 10 µL of a 1:250 dilution of furimazine (Promega) in stimulation buffer was added, and after 120 s, 10 µL of a 10x solution of the compounds in stimulation buffer was added. 180 s after furimazine addition, the plate was placed in a SpectraMax iD5 and read using a program consisting of 60 s of shaking, followed by 500 ms integration reads with a 485/20 nm filter, followed by a 535/25 nm filter.

### Animal ethics

All animal studies were approved by the Institutional Animal Care and Use Committee (IACUC) at the University of Arizona and were conform to the guidelines for the use of laboratory animals of the National Institutes of Health (NIH publication no. 80–23, 1966), or in accordance with the guidelines of the Danish Animal Experimentation Inspectorate (permission number 2021-15-0201-01036) in a fully AAALAC-accredited facility, under the supervision of the local animal welfare committee at the University of Copenhagen.

### Paw incision model of post-operative pain

Male C57Bl/6j mice (6-8 weeks old, The Jackson Laboratory, Bar Harbor, ME) were housed one to six per box (single housing due to fighting) with standard bedding and ad libitum access to standard chow and tap water. Mice were kept in 12 h/12 h light/dark cycles and left to habituate for one week prior to testing. To analyze post-surgery acute incisional hypersensitivity, a plantar incision model was used as described by Brennan *et al*. (*59*). Briefly, a 0.5 cm long incision, from the heel toward the toes, was made through the skin and fascia of the plantar aspect of the left hind paw, including the underlying muscle. The plantaris muscle was then elevated and longitudinally incised, leaving the origin and insertion intact. After hemostasis with gentle pressure, the skin was closed with two mattress sutures of 5-0 nylon on a curved needle.

### Measurement of tactile sensory thresholds for post-operative pain model

Tactile sensory thresholds were assessed by measuring the withdrawal response to probing the plantar surface of the hind paw with a series of calibrated fine filaments (von Frey). Each filament was applied perpendicular to the plantar surface of the paw of mice held in suspended wire mesh cages. The “up and down” method was used to identify the mechanical force required for a paw withdrawal response. The size range of stimuli was between 2.44 (0.4 mN) and 4.56 (39.2 mN). The starting filament was 3.61 (3.9 mN). The filament was placed perpendicular to the skin with a slowly increasing force until it bent; it remained bent for approximately 1 s and was then removed. Data were analyzed using the nonparametric method of Dixon, as described by Chaplan *et al*. (*60*). The results are expressed as the mean withdrawal threshold that induced a paw withdrawal response in 50 % of the animals.

### Spared nerve injury (SNI) model of neuropathic pain

C57BL/6Nrj male mice, approximately 10 weeks of age (Janvier, France), were housed 1–6 per box (single housing due to fighting) with standard bedding and ad libitum access to standard chow (Altromin 1342, Brogaarden, Denmark) and tap water. Mice were kept in 12 h/12 h light/dark cycles (experiments performed during their inactive phase) and left to habituate to the facility for one week prior to testing. Mice were anesthetized with 2 % isoflurane gas followed by subcutaneous injection of buprenorphine (0.1 mg/kg, Department of Experimental Medicine, University of Copenhagen (UCPH)), and SNI surgery was performed on the left hind leg as follows. The skin on the lateral surface was incised for approximately 0.5 cm between the hip and knee, followed by cutaneous application of a single drop of a mixture of 10 mg/kg lidocaine and 5 mg/kg bupivacaine (Department of Experimental Medicine, UCPH) before lengthwise division of the biceps femoris muscle, leading to exposure of the three branches of the sciatic nerve. The sural branch was left intact, whereas the peroneal and tibial branches were ligated with a single surgical knot and axotomized distal to the ligation. Wounds were closed with surgical glue, and animals were monitored daily for signs of stress or discomfort, but in all cases, recovered uneventfully. Immediately post-surgery, the mice were administered carprofen (5 mg/kg, Department of Experimental Medicine, UCPH). On day 16 post-surgery, mice were injected i.p. with a volume of 10 µL/g of the indicated doses of Fj1, 30 mg/kg of gabapentin (Sigma Aldrich), or vehicle (phosphate buffered saline).

### Measurement of tactile sensory thresholds for SNI model

For the SNI experiments, von Frey filaments ranging from 0.04 to 2 g (g = gram-forces) (0.04, 0.07, 0.16, 0.4, 0.6, 1.0, 1.4, 2.0) were used to determine the mechanical paw withdrawal threshold (PWT). Filaments were applied in ascending order to the frontolateral plantar surface of the hind paws innervated by the sural nerve. Mice were placed in red PVC plastic cylinders (8 (Ø) × 7.5 (h) cm) on a wire mesh and allowed to habituate for a minimum of 20 min prior to the experiment initiation. Each von Frey hair was applied five times with adequate resting periods between each application, and the number of withdrawals was recorded. The withdrawal threshold was determined as the von Frey filament eliciting at least three positive trials out of five applications in two consecutive filaments. A positive trial was defined as a sudden paw withdrawal, flinching, and/or paw licking induced by the filament. The animals were habituated to the experimental room for a minimum of 60 min before the initiation of the experiment, and the experimenter (female) was blinded to the treatment groups.

### Structural Preparation

Modelling of consomatin Fj1 at the SSTR_4_ made use of the X-ray crystal structure of consomatin Ro1 (PDB: 7SMU) (*16*), and the cryoEM structure of the SSTR_4_ in complex with the G_i1_ subunit and SST-14 ligand (PDB: 7XMS) (*31*) as modelling templates. Because of the sequence similarity of consomatin ligands and SST-14, specifically the core Trp-Lys receptor-binding motif mimicking the broader conserved Phe-Trp-Lys-Thr residues present within SST ligands, Fj1 was assumed to bind the same orthosteric binding pocket as the endogenous ligands. Modeller (v.10) utility was used to create complexes of ligands bound to the SSTR_4_ binding pocket, generating 50 models with the top model assessed using the Modeller objective function to seed subsequent simulations. Due to the absence of N-terminal SSTR_4_ residues present within the cryoEM structure, only residues Met47-Phe322 were included within the final complex, with the G_i1_ subunit removed. Non-standard amino acids hydroxyproline (Hyp9) and D-tryptophan (D-Trp4) were included, with parameters assigned from the CHARMM36m force field. The resulting complex was inserted into a model POPC membrane using the CHARMM-GUI server and solvated in a box of CHARMM-modified TIP3P (mTIP3P) water, extending > 10 Å beyond all protein atoms. The resulting bilayer spanned an area of 100 × 100 Å^2^, with a total of 130 lipids each in the upper and lower leaflets. Sodium and chloride ions were included to neutralize the system and attain an ionic strength of 0.1 M.

### Molecular Dynamics Simulations

Simulations were performed using the GROMACS v2022) (*61*) package with the CHARMM36m force field (*62, 63*). Temperature coupling was achieved using velocity rescaling applying a coupling time of 0.1 ps with the protein, membrane and water/ions coupled separately at 310 K. Semi-isotropic pressure coupling was maintained using the Parrinello-Rahman method with a coupling time of 2.0 ps. All simulations were performed with a single non-bonded cut-off of 12 Å, with van der Waals interactions switched at 1.0 Å. The Verlet neighbor searching cut-off scheme was applied with a neighbor-list update frequency of 25 steps (50 fs); the time step used in all the simulations was 2 fs. Periodic boundary conditions were applied using the particle-mesh Ewald method to account for long-range electrostatic interactions. The bond lengths were constrained using the P-LINCS algorithm (*64*). Initial minimization and equilibration of all simulations followed established procedures outlined by the CHARMM-GUI via a steepest descent protocol, followed by short positionally restrained equilibration in the NVT (canonical) and NPT ensemble. The systems were then allowed to progress for 1 μs, with each system simulated in 10 replicas, for a total simulation time of 10 μs per consomatin analog.

### Statistical analysis

All statistical analyses, fittings, and plotting of curves were performed using GraphPad Prism 10. For the paw incision model, the effects at different time points were analyzed using two-way ANOVA with Geisser-Greenhouse correction and Dunnett’s post-test for multiple comparisons (all compared to vehicle), while the area under the curve plot was analyzed using one-way ANOVA with Dunnett’s post-test for multiple comparisons (all compared to vehicle). For the SNI model, the effect over time was analyzed as described for the paw incision model (all compared to the vehicle on the ipsilateral side), while the area under the curve was analyzed using a two-way ANOVA with Dunnett’s post-test for multiple comparisons (all compared to the vehicle on the ipsilateral side).

## Supporting information

Supplemental File 2 (Jupyter Notebook)

## List of Supplementary Materials

**Supplemental file 1**

Methods

Figs. S1 to S3

References (*65–70*)

**Supplemental file 2**

Code to extract parameters for PCA.

## Acknowledgments

Prof. Hans Bräuner-Osborne for help with establishing GPCR receptor assays. Dr. Joanna Gajewiak for illustrations of the chemical structure of consomatin Fj1. Dr. Paula Flórez Salcedo for illustration of the cone snail shell.

## Funding

Villum Foundation Young Investigator grant 19063 (HS-H)

Lundbeck Foundation Experiment grant R400-2022-509 to WEB)

Independent Research Fund Denmark grant 3102-00006 (TLK)

Lundbeck Foundation Ascending Investigator grant R344-2020-1063 (KLM)

Lundbeck Postdoc grant R322-2019-1816 (KLJ)

National Institutes of Health R01NS116694 (AP)

National Institutes of Health K08NS104272 (AP)

Part of this work was undertaken with the assistance of resources from the National Computational Infrastructure (NCI), which is supported by the Australian Government and provided through Intersect Australia Ltd.

## Author contributions

Conceptualization: WEB, HS-H

Methodology: WEB, IBLR, TLK, EE, HYY, CMG, KLJ, NAS, LFM

Software: WEB, TLK, NAS

Formal Analysis: WEB, TLK, NAS

Resources, The NCI (supported by the Australian Government and provided through Intersect Australia Ltd)

Investigation: WEB, IBLR, TLK, EE, HYY, CMG, KLJ, NAS, and LFM

Data curation: WEB

Writing – Original Draft: WEB, HS-H

Writing – Review and Editing: WEB, IBLR, TLK, EE, HYY, KLJ, NAS, LFM, BJS, KLM, AP, HS-H

Visualization: WEB, TLK, HS-H

Supervision: WEB, KKS, BJS, KLM, AP, HS-H

Project Administration: WEB and HS

Funding Acquisition: WEB, TLK, KLJ, KLM, HS-H, AP

## Data and materials availability

All data and code can be found either in databases as described in the text, or in the main text of supplemental materials. All materials are commercially available.

