## Supplemental File 1 (Methods and data) for "Venom-inspired somatostatin receptor 4 (SSTR4) agonists as new drug leads for peripheral pain conditions"

#### Supplemental information

**The pdf includes:**
Material and Methods
Figs. S1 to S3
References

**Other Supplementary Material for this manuscript includes the following:**
Jupyter notebook with code for extracting parameters for PCA (Supplemental File 2).

##### Materials and Methods

**Toxin sequence extraction and principal component analysis (PCA)**
Consomatin sequences were extracted from the SRA datasets of cone snails from Bioproject PRNJ526781, as described previously *(17)*. Briefly, data were downloaded from NCBI and preprocessed using fastp v0.20.1 *(65)*. Consomatin-encoding reads were extracted using tblastn with several previously identified consomatins as queries *(16)*. These hits were assembled using Trinity v.2.13.2 *(66, 67)* as single reads. The generated contigs with more than 5-fold coverage were translated in all reading frames and putative toxins were extracted with regular expressions ("[^C]{0,2}C[^C]{0,3}WK[^C]{0,3}C[^RKC]{0,6}") which allows the "WK" motif to appear at any position in the disulfide loop.
We next extracted the length, molecular weight, isoelectric point, hydropathy, aliphaticity, aromaticity, polarity, positive and negative charge, and percentage of unique amino acids (glycine and proline), as well as the count of the 140 most common amino acid 1-to-3-mers across all sequences using an in-house script. Principal components were calculated using sklearn v.1.1.3 *(68)* in Python 3.9.16 on data scaled to remove the mean and unit variance. The first two principal components were visualized in R v.4.2.2 using ggplot v.3.4.2 *(69)*.

**Peptide synthesis and purification**
Peptides were either purchased from Genscript (to > 95 % purity) or synthesized in-house as described below. Peptides were synthesized by solid-phase peptide synthesis (SPPS) at a 0.1 or 0.05 mmol scale using preloaded Fmoc-protected Tentagel R HMPA resin from Rapp Polymere (Tübingen, Germany) for C-terminal carboxylic acid sequences and TentaGel® S RAM Resins for C-terminal amide sequences. Fmoc-protected amino acids, coupling reagents, and solvents used for synthesis were purchased from Iris Biotech (Marktredwitz, Germany). Synthesis was performed using a Syro I instrument (Biotage, Uppsala, Sweden). The coupling conditions were room temperature (RT) for 2 × 120 min using 5.2 equivalents of amino acids, 4.7 equivalents of N-[(1H-benzotriazol-1-yl)(dimethylamino) methylene]-N-methylmethanaminium hexafluorophosphate N-oxide, 5.2 equivalents of 1-hydroxy-7-azabenzotriazol, and 8 equivalents of N,N-diisopropylethylamine in dimethylformamide (DMF) relative to the resin. Deprotection was performed at RT with 40 % piperidine in DMF for 3 min, followed by 20 % piperidine in DMF for 15 min. The washing steps were performed with 2 × N-methyl-2-pyrrolidone, 1 × dichloromethane (DCM), and 1 × DMF. After completion of peptide assembly, the peptidyl resin was washed with 3 × DCM and dried. The peptide was released from approximately 0.025 mmol of resin using a mixture (2 mL) of 95 % trifluoroacetic acid (TFA), 2.5 % triethylsilyl, and 2.5 % water over 2.5 h. For methionine-containing sequences, the 2 mL mixture contained 87.5 % TFA, 2.5 % triethylsilyl, 2.5 % water, 2.5 % thioanisole, and 5 % ethanethiol, or alternatively 87.5 % TFA, 2.5 % triethylsilyl, 2.5 % water, 2.5 % thioanisole, and 5 % ethanethiol with approximately 0.4 mg sodium iodide (NaI). Cold diethyl ether (13 mL stored at -20 °C) was added, and the mixture was further

cooled to -85 °C for 30 min before the peptide was isolated by centrifugation. The peptide was redissolved in a mixture of acetic acid, water, and acetonitrile (ACN) in a volume ratio of 1:10:4 and freeze-dried to remove unwanted carboxylation of Trp. For peptides containing methionine, acetic acid was replaced with half the volume of TFA. The freeze-dried compound was dissolved in a few drops of TFA and 5 mL of ACN, followed by the addition of 5 mL of water. Phosphate buffer (0.1 M) was added to this mixture until a total volume of 18 mL. The pH was adjusted to 6 by using 1 M hydrochloride acid. Dimethyl sulfoxide (2.0 mL) was added (10 %, v/v) and the mixture was stirred for 40–72 h at RT to allow disulfide bond formation. The mixture was filtered and directly purified using one of two methods: the peptides were purified using a Luna C18(2) column from Phenomenex (Torrance, USA, 5 µm, 100 Å, 250 × 10 mm) on a Dionex Ultimate 3000 HPLC system (Thermo Fisher, Waltham, USA) or a 50 g Biotage Sfär Bio C18 D column on a Selekt System (Biotage, Sweden). For both systems, a gradient of 5–100 % ACN in water / 0.1 % formic acid (or 0.1 % TFA for methionine-containing sequences) was applied. The product was isolated and lyophilized by freeze-drying. The final product was analyzed by liquid chromatography-mass spectrometry using a Dionex Ultimate 3000 ultrahigh-performance liquid chromatography system (Thermo Fisher) connected to an Impact HD mass spectrometer (Bruker, Bremen, Germany).

**PRESTO-Tango beta arrestin recruitment assay**

HTLA cells (a kind gift from Prof. Hans Bräuner-Osborne) maintained in DMEM (Gibco) supplemented with 10 % FBS (Biowest, Nuaillé, France), 100 U/mL penicillin / 100 mg/mL streptomycin (Gibco), 100 µg/mL hygromycin B (ThermoFisher Scientific), and 2 µg/mL puromycin (Gibco) (growth medium) were maintained in a water jacketed 5 % $CO_2$ incubator and passaged every 2-3 days using trypsin/EDTA (Gibco). 1E6 cells were seeded into 6 well plates. The following day, the medium was changed, and the cells were transfected with 2 µg of DNA from the relevant receptor construct (obtained from Addgene) and 20 µL PolyFect (Qiagen) into 100 µL DMEM and incubated for 10 min. Growth medium (400 µL) was added and the mixture was added to the cells. 14-18 h later, 4E4 cells per well in 40 µL of DMEM supplemented with 1 % dialyzed FBS (assay medium) were seeded into white clear bottom 386 well plates (Corning). The following day, the medium was changed to 40 µL new assay meduim and compounds at 5x final concentration were added in HBSS supplemented with 20 mM HEPES, 1 mM $CaCl_2$, 1 mM $mgCl_2$, and 0.1 % BSA (stimulation buffer). The following day, the medium was discarded, and 20 µL of a 20x dilution of BrightGlo (Promega) in stimulation buffer was added. The plate was incubated in the dark at RT for 20 min, and luminescence was read at 1000 ms integration time on a SpectraMax iD5 (Molecular Devices).

**Images and visual representation**

All graphs were created using either GraphPad Prism 10 or a custom Python script for heatmaps (https://github.com/Aephir/create_heatmap, v2.0.6 used). All images that are not data graphs or plots were created using Biorender, Adobe Illustrator, and/or Inkscape.

**Data analysis**

The BRET signals in the G protein dissociation assay were calculated by dividing the signal at 535 nm by that at 485 nm. All concentration-response curves were fitted using a variable slope (four-parameter) equation:

$$Y = \frac{R_{min} + (R_{\max} - R_{min})}{1+10^{(logEC_{50}-X)\times Hill\ slope}} \text{ (Equation 1)}$$

Potency calculations were performed using log10 transformed values ($pEC_{50}$) before transforming the average to $EC_{50}$ values.

For the MD data, representative orientations of ligand complexes were determined by clustering all simulation replica trajectories (limited to 100,000 frames) using the GROMOS method with a cut-off between clusters of 0.25 nm. VMD was used along with the tachyon renderer *(70)* for visualization, and the images were encoded to produce videos with ffmpeg.

#### Figure S1

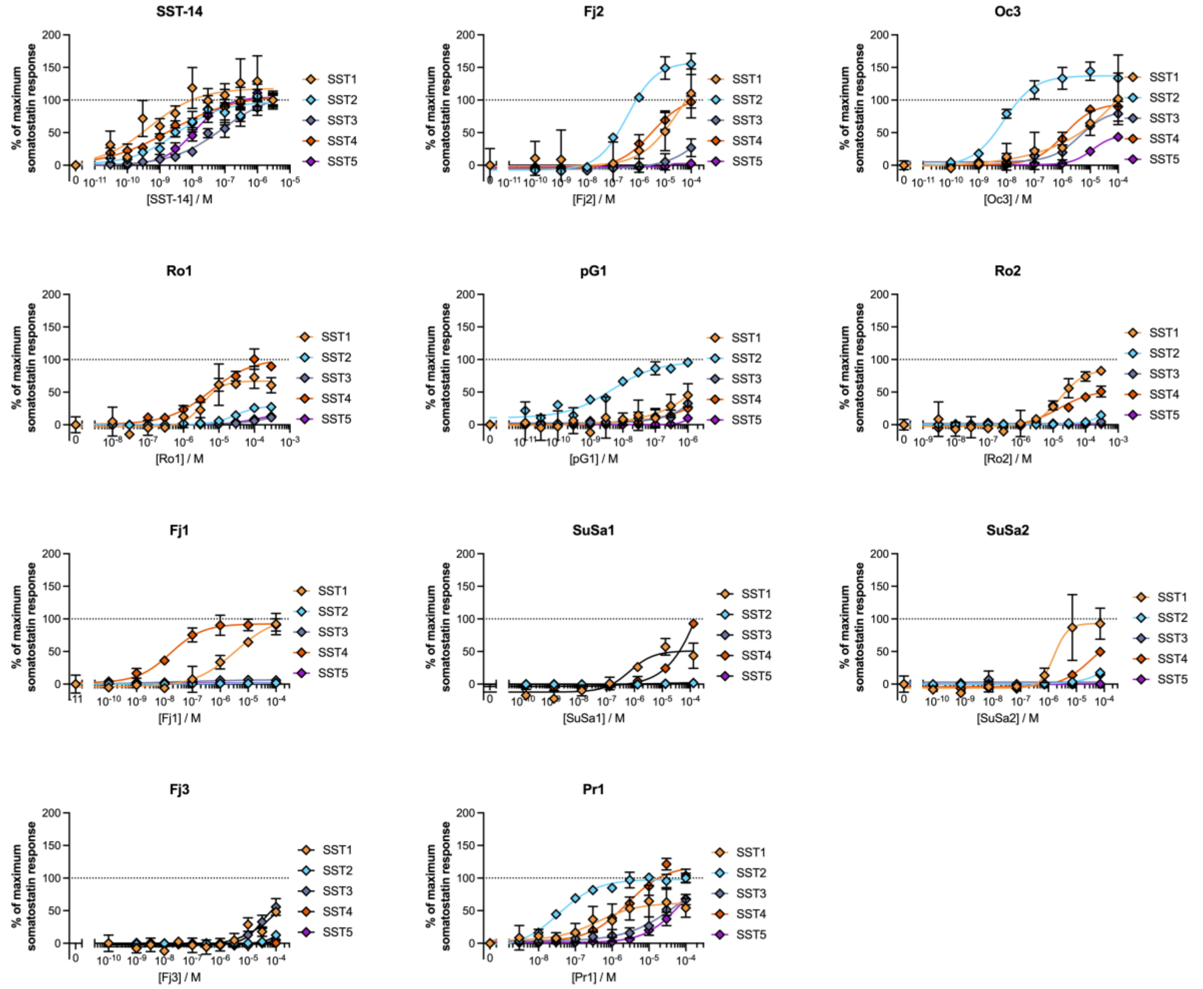


**Fig S1. Representative curves of SST-14 and synthesized consomatins in PRESTO-Tango assay.** Datapoints represent the mean of three technical replicates, and error bars represent standard deviations (related to Fig. 1).

#### Figure S2

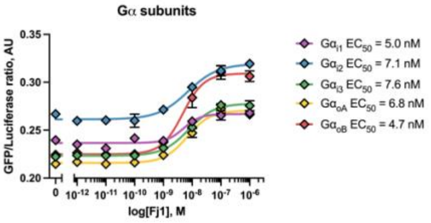


**Fig S2. Effect of Fj1 at different Gα subunits from the G$\alpha_{i/o}$ family.** The concentration-response curve of Fj1 at the SSTR$_4$ using five different Gα subunits from the G$\alpha_{i/o}$ family. Datapoints represent the mean of two technical replicates, and error bars represent standard deviations (related to Fig. 2).

### Figure S3

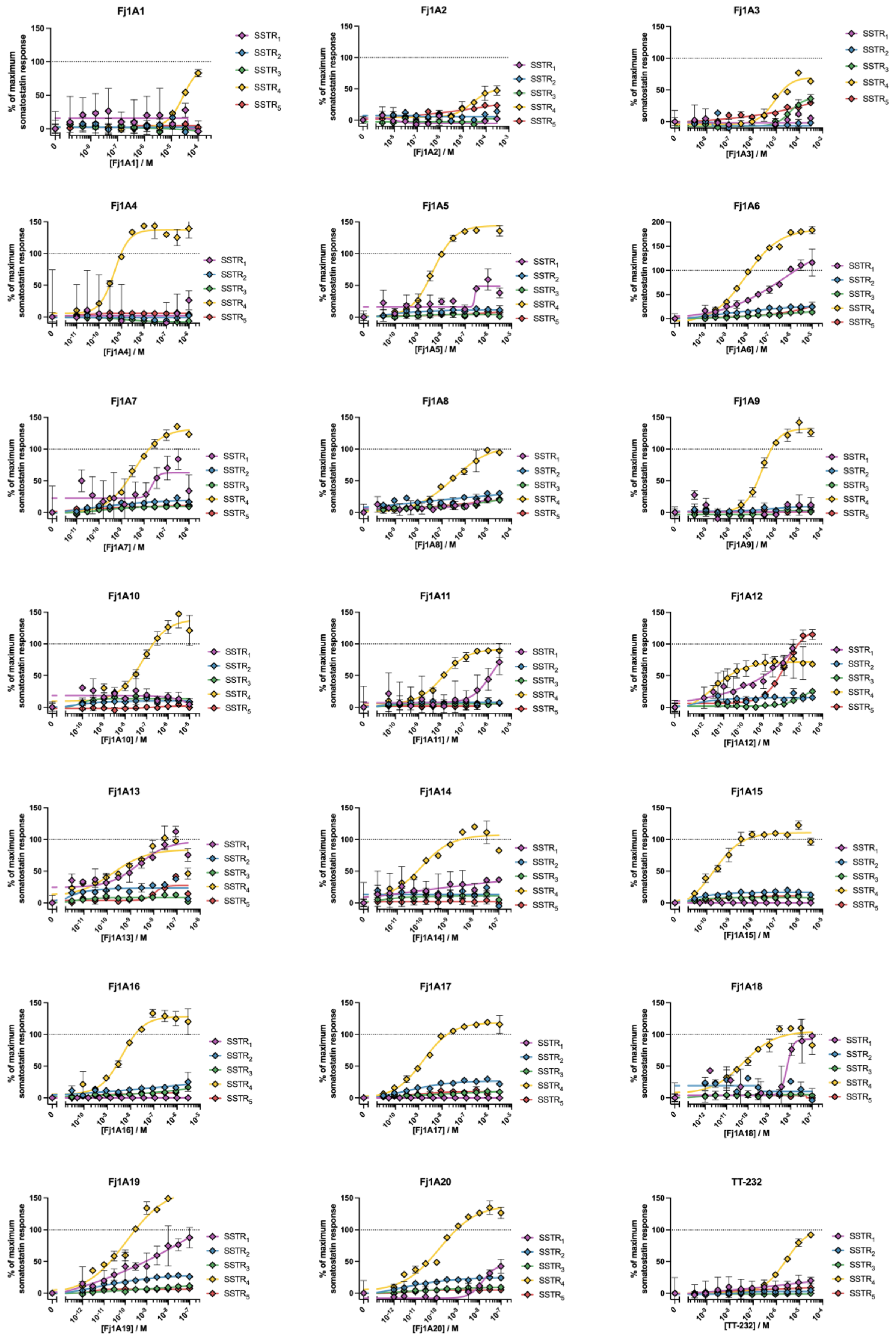
Fj1A1
Fj1A2
Fj1A3
Fj1A4
Fj1A5
Fj1A6
Fj1A7
Fj1A8
Fj1A9
Fj1A10
Fj1A11
Fj1A12
Fj1A13
Fj1A14
Fj1A15
Fj1A16
Fj1A17
Fj1A18
Fj1A19
Fj1A20
TT-232
% of maximum somatostatin response
[Fj1A1] / M
[Fj1A2] / M
[Fj1A3] / M
[Fj1A4] / M
[Fj1A5] / M
[Fj1A6] / M
[Fj1A7] / M
[Fj1A8] / M
[Fj1A9] / M
[Fj1A10] / M
[Fj1A11] / M
[Fj1A12] / M
[Fj1A13] / M
[Fj1A14] / M
[Fj1A15] / M
[Fj1A16] / M
[Fj1A17] / M
[Fj1A18] / M
[Fj1A19] / M
[Fj1A20] / M
[TT-232] / M
$SSTR_1$
$SSTR_2$
$SSTR_3$
$SSTR_4$
$SSTR_5$

**Fig S3. Representative curves of Fj1 analogs in G protein dissociation assay.** Datapoints represent the mean of two technical replicates, and error bars represent standard deviations (related to Fig. 5).
