## Supplemental File 2 (Jupyter Notebook) for "Venom-inspired somatostatin receptor 4 (SSTR4) agonists as new drug leads for peripheral pain conditions": Supplemental File 2.ipynb

```
# %% [code]
from Bio import SeqIO
from Bio.SeqUtils.ProtParam import ProteinAnalysis
from Bio.SeqIO.FastaIO import SimpleFastaParser 
import pandas as pd
import numpy as np
from sklearn.decomposition import PCA
from sklearn.preprocessing import StandardScaler
import re
from sklearn.feature_extraction.text import CountVectorizer
import matplotlib.pyplot as plt
from sklearn.manifold import MDS

# %% [code]
def get_bio(seq):
    
    with open('Prey.species.Lautoconus_split.txt') as prey_file:
        conus_genus = dict()
        conus_prey = dict()
        conus_strat = dict()
        for line in prey_file:
            line = line.rstrip().split('\t')
            conus_genus[line[0]] = line[1].split()[0]
            conus_prey[line[0]] = line[2]
            conus_strat[line[0]] = line[3]
        
    if 'XXA' in seq:
        return [seq,'Annelid','Annelid']
    if 'XXM' in seq:
        return [seq,'Mollusk','Mollusk']
    if 'XXV' in seq:
        return [seq,'Vertebrate','Vertebrate']
    else:
        return [seq,conus_prey[seq],conus_prey[seq]]


# %% [code]
# Used for figure. Do not change code to play around
def read_fasta(file_path) :
    with open(file_path) as fasta_file :
        records = {}
        corpus = []
        titles = []
        nr = 0
        for title, sequence in SimpleFastaParser(fasta_file):
            record = get_bio(title)
            title = f'{title}_{nr}'
            nr += 1
            
            record.append(sequence) #sequence
            record.append(len(sequence)) #sequence length
            
            analysed_seq = ProteinAnalysis(sequence)
            record.append(analysed_seq.molecular_weight()) #molecular weight
            record.append(analysed_seq.isoelectric_point()) #isoelectric point
            record.append(analysed_seq.gravy())

            record.append((sequence.count('A')+sequence.count('L')+sequence.count('V')+sequence.count('I'))
                          /len(sequence))
            record.append((sequence.count('F')+sequence.count('W')+sequence.count('Y'))
                          /len(sequence))
            record.append((sequence.count('N')+sequence.count('C')+sequence.count('Q')+
                           sequence.count('M')+sequence.count('S')+sequence.count('T'))
                          /len(sequence))
            record.append((sequence.count('G')+sequence.count('P'))
                          /len(sequence))
            record.append((sequence.count('R')+sequence.count('K')+sequence.count('H'))
                          /len(sequence))
            record.append((sequence.count('E')+sequence.count('D'))
                          /len(sequence))
            
            records[title] = record
            
            names = ['species', 'prey', 'type','sequence', 'length', 'mW', 'iep', 'gravy', '%ali', '%aro', '%pol',
                    '%uniq', '%pos', '%neg']
            
            corpus.append(sequence)
            titles.append(title)
    vectorizer = CountVectorizer(analyzer='char', ngram_range=(1, 3), max_features = 140)
    X = vectorizer.fit_transform(corpus)
    
    aa_features_df = pd.DataFrame(X.toarray(), columns=vectorizer.get_feature_names_out(), index = titles)
    phychem_features_df = pd.DataFrame(records, index = names)
    
    x = pd.concat([phychem_features_df.T, aa_features_df], axis = 1)
            
    return x

# %% [code]
def run(infile, outfile):
    tox_df = read_fasta(infile)
    tox_df_red = tox_df.drop(['species', 'prey', 'type','sequence'], axis = 1)
    
    scaler = StandardScaler()
    scaler.fit(tox_df_red)
    tox_df_red_scale = scaler.transform(tox_df_red)
    
    pca = PCA(n_components=5)
    #fitting = MDS(n_components=2).fit_transform(tox_df_red_scale)
    fitting = pca.fit(tox_df_red_scale).transform(tox_df_red_scale)
    
    q = tox_df
    q["PC1"] = fitting.T[0]
    q["PC2"] = fitting.T[1]
    q["PC3"] = fitting.T[2]
    q["PC4"] = fitting.T[3]
    q["PC5"] = fitting.T[4]
    
    q.to_csv(outfile)
    
    return q

# %% [code]
q = run(path/to/fasta,path/to/outfile)

# %% [code]
q
```
